# Beyond the skin barrier: optical clearing enables non-invasive cortex-wide optical coherence angiography in mice in-vivo

**DOI:** 10.64898/2026.03.02.709062

**Authors:** Daewoon Seong, Sungbin Yun, Sangyeob Han, Sagor Biswas, Bosung Kim, Eva Remlova, Daniel Razansky, Jeehyun Kim, Zihao Ou, Mansik Jeon

## Abstract

Non-invasive, high-resolution visualization of mouse brain vasculature remains challenging due to significant light scattering and absorption by mammalian tissues, hence many optical imaging protocols require scalp and/or skull excision. Here we present a fully reversible tartrazine-based optical clearing strategy that enables cortex-wide optical coherence tomography angiography (OCTA) through intact scalp and skull. We characterized tartrazine properties in the near infrared (NIR)-II band of the 1.3 µm swept-source OCTA system, confirming minimal absorption across 1.25–1.35 µm wavelength range and an effectively constant refractive index, suggesting negligible OCTA distortions. Spatially selective agent application showed that intracranial vessels emerge selectively within the treated region of interest (ROI), whereas untreated regions retain strong interference by the scalp vascular features. Depth-encoded projections and cross-sectional OCTA demonstrated an increased signal recovery at depth and an extended vessel-detection range after clearing. Vessel-map changes were quantified using intersection-over-union and Dice coefficients, yielding high similarity outside the ROI and reduced similarity within the ROI, consistent with a transition from scalp to brain vasculature. Reproducibility was confirmed in three independent 11-week-old mice and validated against scalp-removed reference OCTA. Screening tartrazine in the 0.3–0.8 Molar concentration range (7-min application) identified 0.6 M as optimal for whole-cortex scanning, balancing clearing efficacy and solution handling. Finally, the protocol generalized across mice aged 5–18 weeks. This approach provides a practical route to non-invasive structural cerebrovascular mapping with OCTA.

## Introduction

Non-invasive, high-resolution delineation of cerebral vascular architecture in the intact mouse brain is essential for preclinical brain imaging, providing a structural reference for studies of neurovascular organization and cerebrovascular biology^1-3^. Optical modalities provide a direct route to resolving cerebrovascular morphology in-vivo at micrometer scale resolution, complementing macroscopic angiography by enabling volumetric mapping of pial vessels, penetrating vessels, and capillary beds^4-6^. Two-photon laser scanning microscopy established intravital 3D fluorescence imaging in scattering tissues and remains a core platform for microvascular angiography in the rodent cortex^7,8^. Three-photon microscopy has further expanded transcranial optical access, enabling visualization of cerebral vasculature through the intact adult mouse skull^9,10^. In addition, photoacoustic microscopy offers a complementary, hemoglobin-based contrast mechanism for high-resolution vascular mapping, including transcranial implementations in mice^11,12^. Furthermore, optical coherence tomography angiography (OCTA) provides label-free, depth-resolved vascular imaging, which extracts motion contrast from flowing red blood cells to generate 3D microangiograms well-suited for longitudinal structural quantification^13-16^. However, because the scalp imposes substantial light scattering and absorption, transcranial optical imaging often still relies on surgical scalp excision to expose the skull, making the procedure inherently invasive^17,18^. These considerations motivate imaging strategies that preserve the native scalp–skull barrier while enabling high-resolution visualization of the cerebral vasculature in-vivo.

A long-standing alternative is in-vivo skin optical clearing, which transiently reduces scattering by modulating tissue hydration and refractive-index mismatch within the skin^19,20^. Diverse topical agents, including polyols and sugars (e.g., glycerol, glucose, fructose), as well as higher refractive-index solvents and polymers (e.g., propylene glycol, PEG-400), have been shown to enhance optical transmission and improve imaging performance across optical modalities^21,22^. Because the stratum corneum restricts diffusion, clearing is often strengthened using chemical permeation enhancers (e.g., thiazone, DMSO, propanediol) and physical delivery strategies that transiently modulate the barrier, such as massage, microneedle-generated microchannels, or ultrasound/sonophoresis^23,24^. More recently, an absorbing-molecule paradigm has been introduced in which topical tartrazine reversibly increases transparency of the shaved mouse scalp, enabling visualization of cerebral vessels without surgical scalp excision, consistent with dye-mediated refractive-index modulation described by the Kramers–Kronig relations^25-28^. Tartrazine has also been applied in OCT-based imaging contexts, including ex-vivo porcine eyes^29^ and in-vivo mouse abdominal skin^30-32^. However, these demonstrations were not designed for transcranial imaging and therefore did not address the compounded optical barrier imposed by the intact scalp and skull for in-vivo mouse brain vasculature imaging.

In this paper, we introduced OCTA cerebrovascular imaging through an intact mouse scalp assisted with tartrazine-based optical clearing. To establish feasibility in the spectral regime of our system, we first characterized the optical behavior of tartrazine in the NIR-II window corresponding to the OCTA illumination band. Then, we performed spatially controlled topical application of the agent and compared OCTA before and after clearing, showing that intracranial vasculature becomes visible specifically within the tartrazine treated region, whereas untreated regions predominantly preserve scalp-vascular features. This effect of tartrazine-based optical clearing was quantitatively supported using segmentation overlap metrics, including intersection-over-union (IoU) and the Dice coefficient. We further assessed experimental consistency by repeating the protocol across three mice, and validated anatomical accuracy by directly comparing intact-scalp, clearing-enabled cerebrovascular maps with corresponding OCTA acquired after scalp removal. To optimize the clearing condition, we screened tartrazine solutions in 0.3-0.8 M concentration range in 0.1 M increments under a fixed 7 minutes application time, identifying 0.6 M as an optimal molar concentration for noninvasive OCTA-based whole mouse cortex vasculature imaging. Finally, we extended the protocol to mice of different ages, demonstrating the broader applicability of tartrazine-assisted intact-scalp OCTA for structural cerebrovascular imaging.

## Results

### Optical characteristics of tartrazine at NIR-II window and experiment procedure

As an initial step, we assessed the optical properties of tartrazine in the NIR–II window, which corresponds to the operating band of the OCTA source used in this study. To evaluate the transparency of tartrazine solutions at different molar concentrations, we measured their spectral transmittance using an ultraviolet-visible (UV–VIS)–NIR spectrometer over the relevant wavelength range (Fig. 1a, Methods, and Supplementary Table 1). The measured transmittance confirmed minimal absorption of tartrazine within the OCT source band (1.25–1.35 µm), which demonstrates the feasibility of tartrazine-assisted optical clearing with OCTA imaging technique. Next, we measured the refractive index of the tartrazine solution within the OCT source band to verify the degree of variance affecting the obtained OCT signal. Because the measurable range of the ellipsometer was limited to wavelengths below 1.0 µm (from 0.6 to 1.0 µm) using ellipsometry (Supplementary Fig. 1 and Methods), we extrapolated the refractive index to the NIR–II region using a Cauchy dispersion model (Supplementary Fig. 2 and Methods). The calculated refractive index remained effectively constant across the OCT source wavelength range (1.25–1.35 µm; Fig. 1b). Accordingly, tartrazine is not expected to introduce appreciable wavelength-dependent refractive-index variations within this band, suggesting that OCT images are unlikely to be distorted by refractive-index dispersion after tartrazine application. The overall tartrazine-based intact-scalp mouse whole-cortex OCTA imaging workflow consists of four steps (Fig. 1c and Methods). Following hair removal, baseline OCTA is first performed to image the superficial scalp vasculature. Tartrazine solution is then applied to the scalp, and the tissue is gently massaged for 7 min to induce optical clearing, after which OCTA is performed to visualize the cerebral vasculature. All imaging was conducted using a customized swept-source OCT system with a central wavelength of 1.3 µm (Fig. 1d and Methods).

**Fig. 1.**
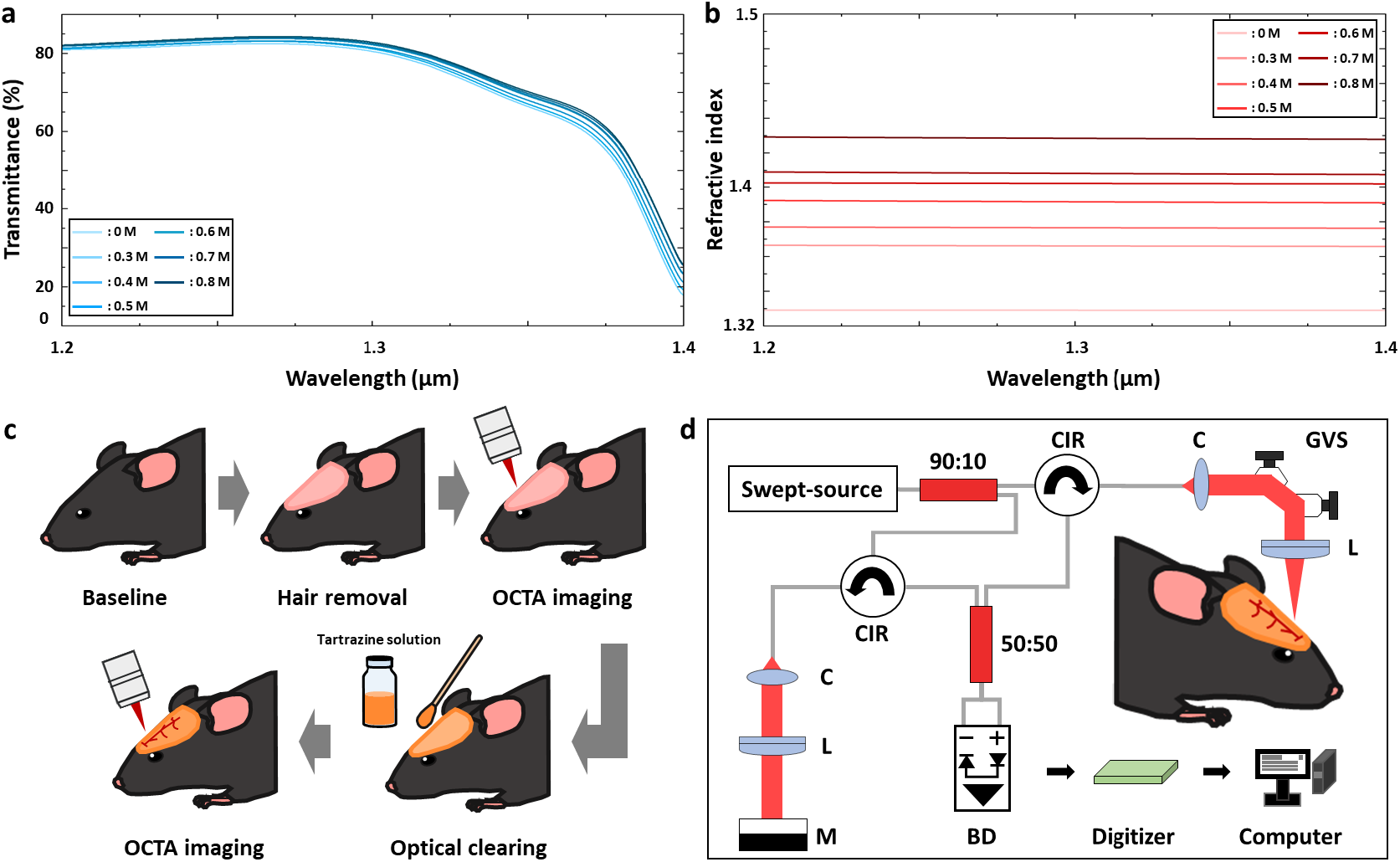
Optical characteristics of tartrazine and overall experimental procedure. **a** Optical transparency of the 1 mm thickness tartrazine solutions with different molar concentrations measured with UV-VIS-NIR spectrometer in the 1.2 to 1.4 µm wavelength range. **b** Cauchy model-based calculated refractive indices of the tartrazine solutions with different molar concentration in the 1.2 to 1.4 µm wavelength range. **c** Overall experimental procedure for tartrazine-assisted intact-scalp whole mouse brain vasculature imaging. **d** Optical configuration of the used OCTA system. BD, balanced detector; C, collimator; CIR, circulator; GVS, galvanometer scanner; L, lens; M, mirror.

### Validation of tartrazine-assisted intact-scalp OCTA for cerebrovascular imaging

To verify the performance of the proposed tartrazine-based intact-scalp whole-brain vasculature imaging using OCTA, we first applied the protocol to an 11-week-old mouse and compared OCTA images acquired before optical clearing and after clearing (Fig. 2a). Before optical clearing, vascular signals were dominated by the superficial scalp vasculature, and limited light penetration through the intact scalp prevented reliable visualization of intracranial vessels. After tartrazine-based optical clearing, reduced scattering within the scalp enabled OCTA visualization of brain vasculature while leaving the scalp intact, a key advantage of the proposed approach. This contrast is more evident in the magnified OCTA views (Fig. 2b and Fig. 2c). In the region without tartrazine application (i.e., outside the region of interest, ROI; yellow-dotted square in Fig. 2a), scalp vessels remain prominently visible in both before and after clearing images, indicating that superficial scalp vascular features are largely preserved (Fig. 2b). In contrast, within the tartrazine-treated ROI (green-dashed square in Fig. 2a), after clearing OCTA reveals vascular features that are clearly distinct from the before clearing superficial scalp vasculature and are consistent with intracranial vessels, indicating that tartrazine increases scalp transparency to enable brain vasculature imaging (Fig. 2c). The tartrazine-treated region, which becomes visibly transparent, is also apparent in the white-light photographs acquired before and after optical clearing (Fig. 2d) and in the OCT maximum intensity projection (MIP) images (Fig. 2e). Consistent with improved optical access, depth-encoded (color-mapped) vascular images show that signals are concentrated near the surface before clearing, whereas vessels at comparatively greater depths become visible after clearing (Fig. 2f). This enhanced penetration is further supported by cross-sectional OCT (Fig. 2g) and OCTA (Fig. 2h) images extracted along the red-dashed line in Fig. 2a, which show increased OCT signal intensity at depth and an extended depth range of vessel detection after clearing. Finally, we quantified the clearing-induced changes by computing IoU (Fig. 2i) and Dice coefficient (Fig. 2j) between before and after clearing images in the out of ROI and ROI regions; as expected, the out of ROI comparisons yielded higher IoU and Dice values than the ROI comparisons because scalp vessels are preserved outside the ROI, whereas the ROI transitions from predominantly scalp vasculature to intracranial vasculature after clearing. Based on these results, the feasibility of tartrazine-enabled intact-scalp OCTA for whole-brain vasculature imaging in mice is verified.

**Fig. 2.**
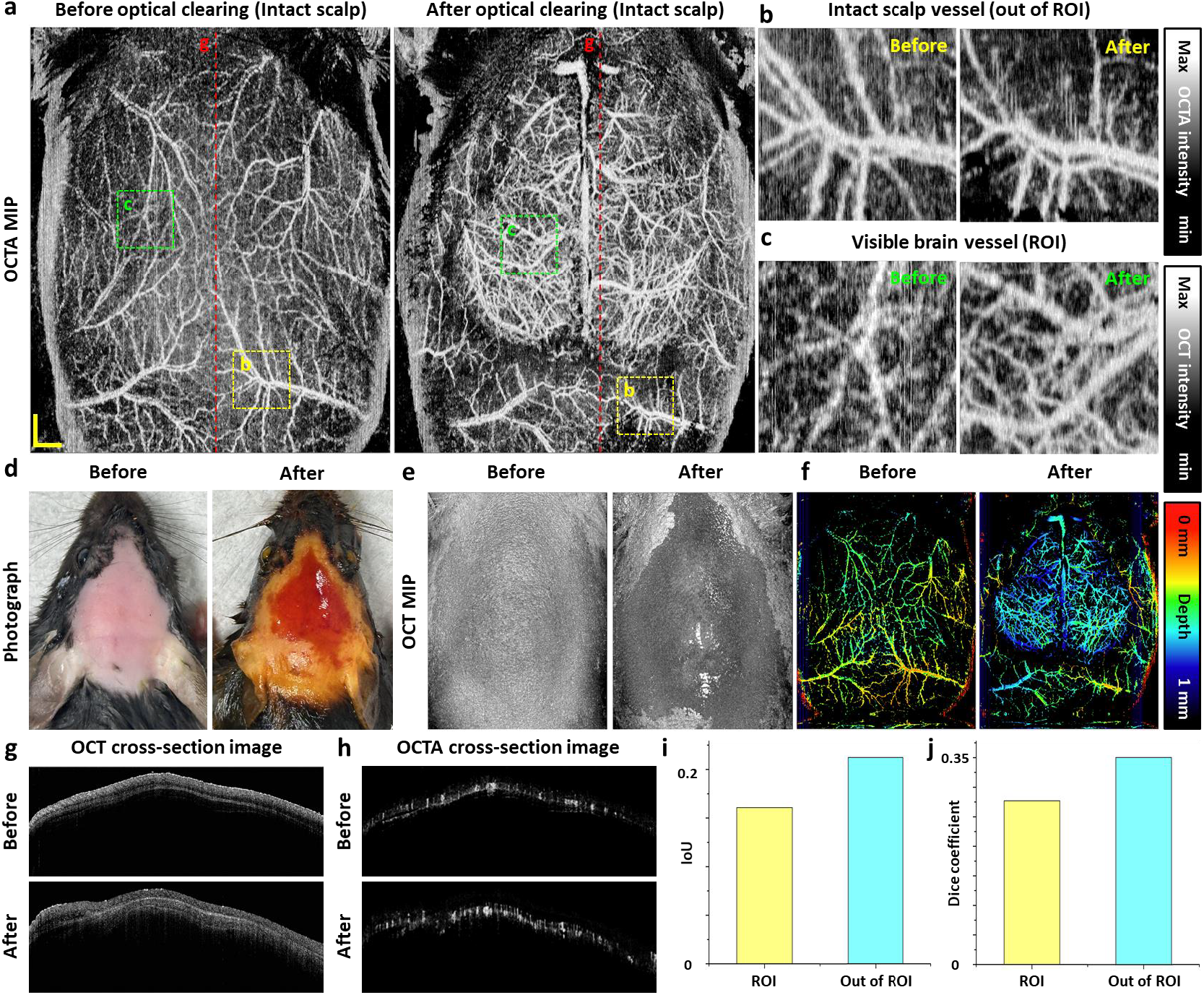
Quantitative results comparing brain vessel images acquired before and after tartrazine-based optical clearing. **a** OCTA MIP images of intact-scalp whole mouse brain vasculature at before and after optical clearing. **b** Magnified OCTA MIP images of out of ROI (i.e., without clearing region) indicated with yellow-dotted square at **a. c** Magnified OCTA MIP images of ROI (i.e., optical clearing region) indicated with green-dotted square at **a. d** Photographs of before and after optical clearing. **e** OCT MIP images of before and after optical clearing corresponding to **d. f** Depth encoded map images of before and after optical clearing to identify the depth positions of each vessel. **g** Representative cross-sectional OCT images of before and after optical clearing extracted from red-dotted line at **a. h** Representative cross-sectional OCTA images of before and after optical clearing extracted from red-dotted line at **a. i** Calculated IoU values using magnified images at out of ROI (**b**) and ROI (**c**). **j** Calculated Dice coefficient values using magnified images at out of ROI (**b**) and ROI (**c**). IoU, intersection over union; OCT, optical coherence tomography; OCTA, optical coherence tomography angiography; ROI, region of interest.

### Reproducibility of the proposed tartrazine-assisted intact-scalp OCTA imaging protocol

Next, to evaluate the repeatability and reproducibility of the proposed method, we applied the identical tartrazine-based optical clearing and intact-scalp OCTA acquisition protocol in three independent 11-week-old mice (Fig. 3a). In all animals, OCTA MIP images consistently showed that optical clearing enabled visualization of intracranial vasculature compared with the pre-clearing condition. Depth-encoded (color-mapped) MIP images further show that, before clearing, the signal is dominated by superficial scalp vessels, whereas after clearing, vessels at comparatively greater depths become visible (Supplementary Fig. 3). To verify that the vessels observed after clearing originated from the brain rather than the scalp, we acquired OCTA images after surgical scalp removal and compared them with after clearing intact-scalp OCTA images. This comparison verified that the vascular networks observed after clearing were consistent with the mouse brain vasculature. This trend is more apparent in the magnified views. In the region indicated by the yellow-dashed box in Fig. 3a, comparisons between before and after clearing images (Fig. 3b, Fig. 3d, and Fig. 3f) exhibited markedly different vascular morphologies, consistent with a transition from superficial scalp vessels to intracranial vessels after clearing. In contrast, comparisons between post-clearing intact-scalp images and the images acquired after scalp removal (Fig. 3c, Fig. 3e, and Fig. 3g) reveal highly similar vascular patterns, as both conditions facilitate visualization of cerebral vessels. These observations provide further evidence of the optical clearing effect and support the cerebral origin of the vasculature visualized after tartrazine treatment. Additionally, the line intensity profile extracted from Fig. 2b further supports that the vascular patterns differ between before and after clearing conditions (Fig. 2h). In contrast, the line intensity profile extracted from Fig. 2c shows close agreement between after clearing intact-scalp OCTA image and the OCTA image acquired after scalp removal, corroborating that the vessels visualized after optical clearing originate from the brain rather than the scalp. Finally, it is quantitatively supported by IoU and Dice coefficient values calculated from the magnified images (Fig. 2b-2g), which demonstrate substantially higher similarity between after optical clearing and scalp removed images than between before and after optical clearing conditions. Together, these results demonstrate the repeatability and reproducibility of the proposed method and confirm that the vasculature visualized after clearing corresponds to cerebral vessels.

**Fig. 3.**
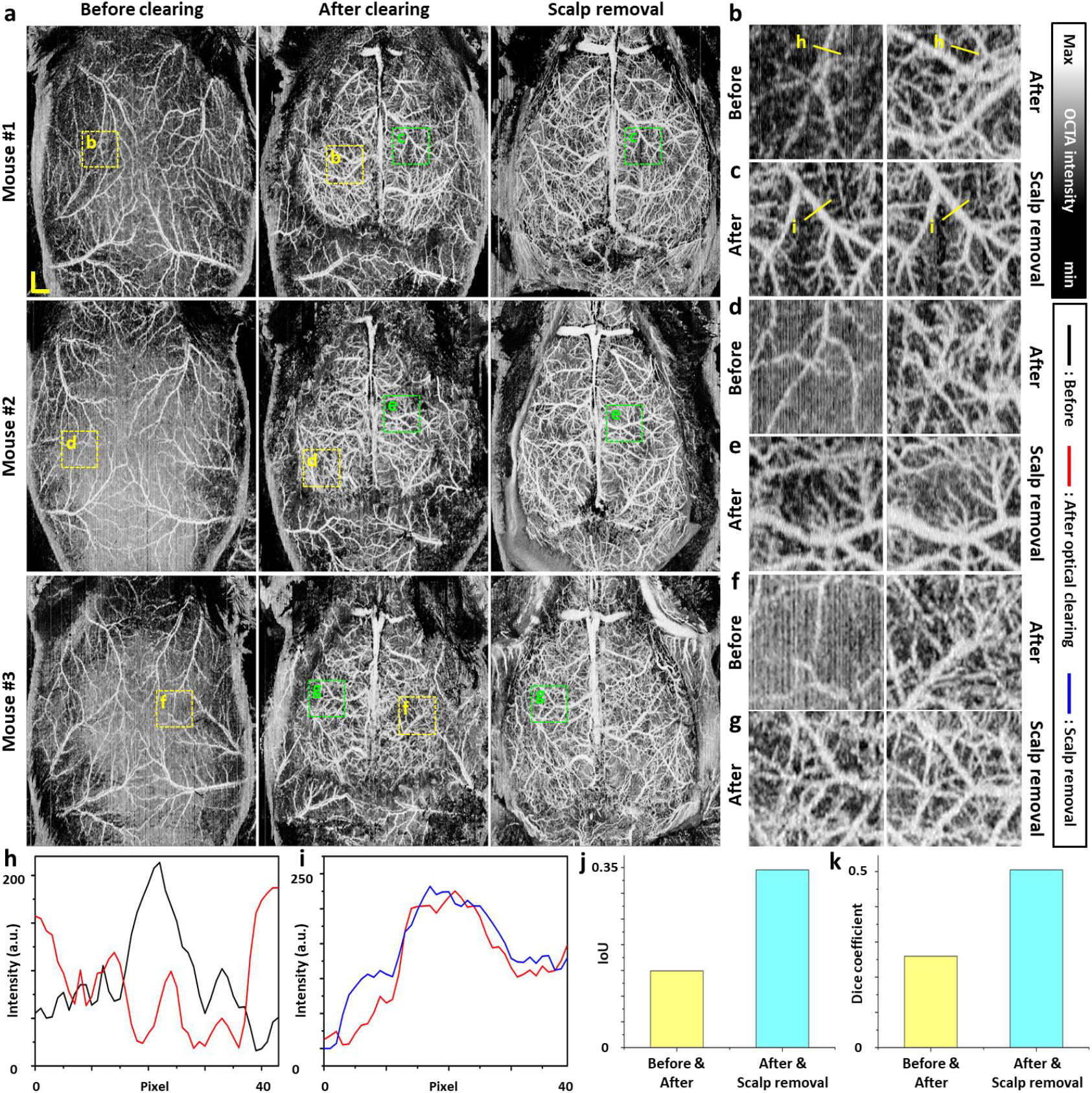
Repeated verification results of tartrazine-based optical clearing for intact-scalp mouse whole brain imaging. **a** OCTA MIP images of three-difference mice acquired at before optical clearing, after optical clearing, and scalp removal. **b, d, f** Magnified OCTA images of before and after optical clearing of mouse #1∼#3 indicated with yellow-dashed square at **a. c** Magnified OCTA images of after optical clearing of mouse #1 indicated with yellow-dashed square at **a. c, e, g** Magnified OCTA images of after optical clearing and scalp removal of mouse #1∼#3 indicated with green-dashed square at **a. h, i** OCTA intensity profile extracted from yellow line at **b, c. j** Calculated IoU values using magnified images at before/after optical clearing and after optical clearing/scalp removal. **k** Calculated Dice coefficient values using magnified images at before/after optical clearing and after optical clearing/scalp removal. IoU, intersection over union; OCTA, optical coherence tomography angiography. Scale bar: 1 mm.

### Optimization of absorbing molecule concentration for intact-scalp OCTA imaging

Next, to optimize the absorbing-molecule–based protocol for intact-scalp OCTA imaging of mouse brain vasculature, we assessed optical clearing efficacy as a function of tartrazine molar concentration. For an objective comparison, we prepared tartrazine solutions from 0.3 M to 0.8 M in 0.1 M increments and applied a standardized protocol across all conditions that consisted of 7 min of massage-assisted topical application followed by OCTA imaging (Supplementary Table 1 and Methods). Compared with before clearing condition, little to no clearing was observed at 0.3 M and 0.4 M (Fig. 4a). Scalp transparency increased progressively with concentration, and robust clearing was consistently achieved from 0.5 M onward (Fig. 4a). Consistent with improved optical access, depth-encoded MIP images showed that vascular signals were concentrated near the surface before clearing and became detectable at greater depths after clearing (Supplementary Fig. 4). The cerebral origin of vessels visualized after clearing was further confirmed by comparison with OCTA acquired after scalp removal. To independently visualize concentration-dependent differences in transmission, we removed the cleared scalp tissue, placed it over the letters “KNU,” and acquired photographs that directly reflected the degree of transparency across conditions (Fig. 4b). The letters were clearly visible at concentrations of 0.5 M and above, but remained poorly discernible at 0.3 M and 0.4 M, consistent with the in-vivo OCTA observations. For quantitative assessment, we extracted line intensity profiles along the yellow line in Fig. 4b. The 0.3 M and 0.4 M conditions exhibited shallow intensity gradients, indicating insufficient optical transmission, and confirming that these concentrations were inadequate for effective scalp clearing. In addition, IoU and Dice coefficients computed from magnified OCTA images (Supplementary Fig. 5) showed high similarity between the after-clearing and scalp-removed conditions at concentrations of 0.5 M and above (Fig. 4d and Fig. 4e). Although clearing was also effective at 0.7 M and 0.8 M, these higher-concentration solutions tended to solidify rapidly over time, limiting their applicability for acquiring a whole-brain OCTA image. This limitation is evident in time-lapse photographs acquired after dispensing a single drop of solution at each concentration and in whole-brain OCTA images in which regions obscured by solidified solution during scanning are marked with yellow circles (Supplementary Fig. 6 and Supplementary Fig. 7). Moreover, under the equal protocol including massaging time, 0.5 M produced a smaller optically cleared area than 0.6 M. Taken together with the rapid solidification at higher concentrations, these results identify 0.6 M as an optimal concentration that provides a sufficiently large cleared field while remaining practical for whole-brain OCTA acquisition within the available time window.

**Fig. 4.**
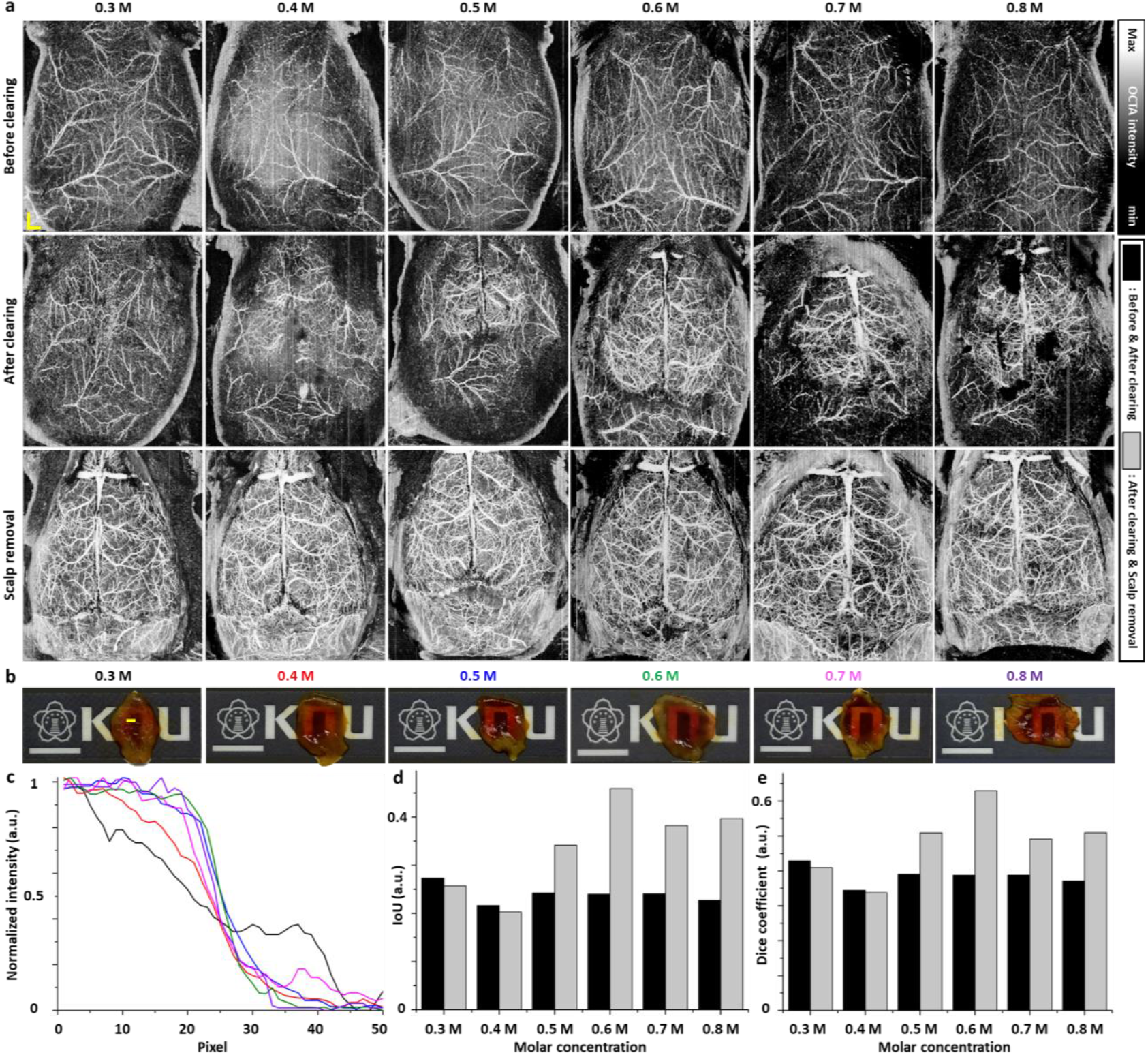
Experimental results for selecting the optimal molar concentration from 0.3 M to 0.8 M. **a** OCTA MIP images of different molar concentrations acquired at before optical clearing, after optical clearing, and scalp removal. **b** Photographs illustrating the difference in the transparency of optically cleared mouse scalp with an increasing concentration. **c** Line profile of mouse scalp indicated with yellow line at **b**. The plot line colors correspond to concentrations colored in panel **b. d** Calculated IoU values of different molar concentrations using magnified images at before/after optical clearing and after optical clearing/scalp removal. **e** Calculated Dice coefficient values of different molar concentrations using magnified images at before/after optical clearing and after optical clearing/scalp removal. IoU, intersection over union; OCTA, optical coherence tomography angiography. Scale bar at **a**: 1 mm. Scale bar at **b**: 1 cm.

### Applicability of tartrazine-enabled intact-scalp OCTA across age groups

To assess the applicability of the proposed tartrazine-based intact-scalp whole-brain OCTA imaging protocol, we tested it in mice aged 5–18 weeks. Compared with before clearing OCTA MIP images, tartrazine-based optical clearing was consistently achieved across all age groups (Fig. 5a). Comparison with OCTA MIP images acquired after scalp removal further confirmed that the vasculature visualized after clearing corresponded to cerebral vessels. In the corresponding depth-coded maps, tartrazine-based clearing also revealed vessels at depths beyond the scalp, supporting visualization of intracranial vasculature (Supplementary Fig. 8). To provide an independent visual assessment of transparency, we excised an untreated scalp sample and a tartrazine-treated, optically cleared scalp sample, placed each over the letters “KNU,” and acquired photographs (Fig. 5b). These photographs indicate that scalp thickness varies across age groups. Line intensity profiles extracted along the yellow line in before clearing images of Fig. 5b showed that the intensity gradient decreased with increasing age (Fig. 5c). In contrast, line intensity profiles extracted from after clearing images of Fig. 5b exhibited steeper gradients than those from before clearing condition across all age groups. The averaged gradient increased to 5.8 after clearing, compared with 3.7 before clearing, further supporting the optical clearing effect. Finally, IoU and Dice coefficient values computed from magnified OCTA images (Supplementary Fig. 9) provided quantitative support for these observations, showing consistently higher similarity across age groups than the corresponding reference values from before versus after clearing comparisons. These results demonstrate that the proposed tartrazine-based intact-scalp OCTA brain imaging protocol is applicable across the tested age range.

**Fig. 5.**
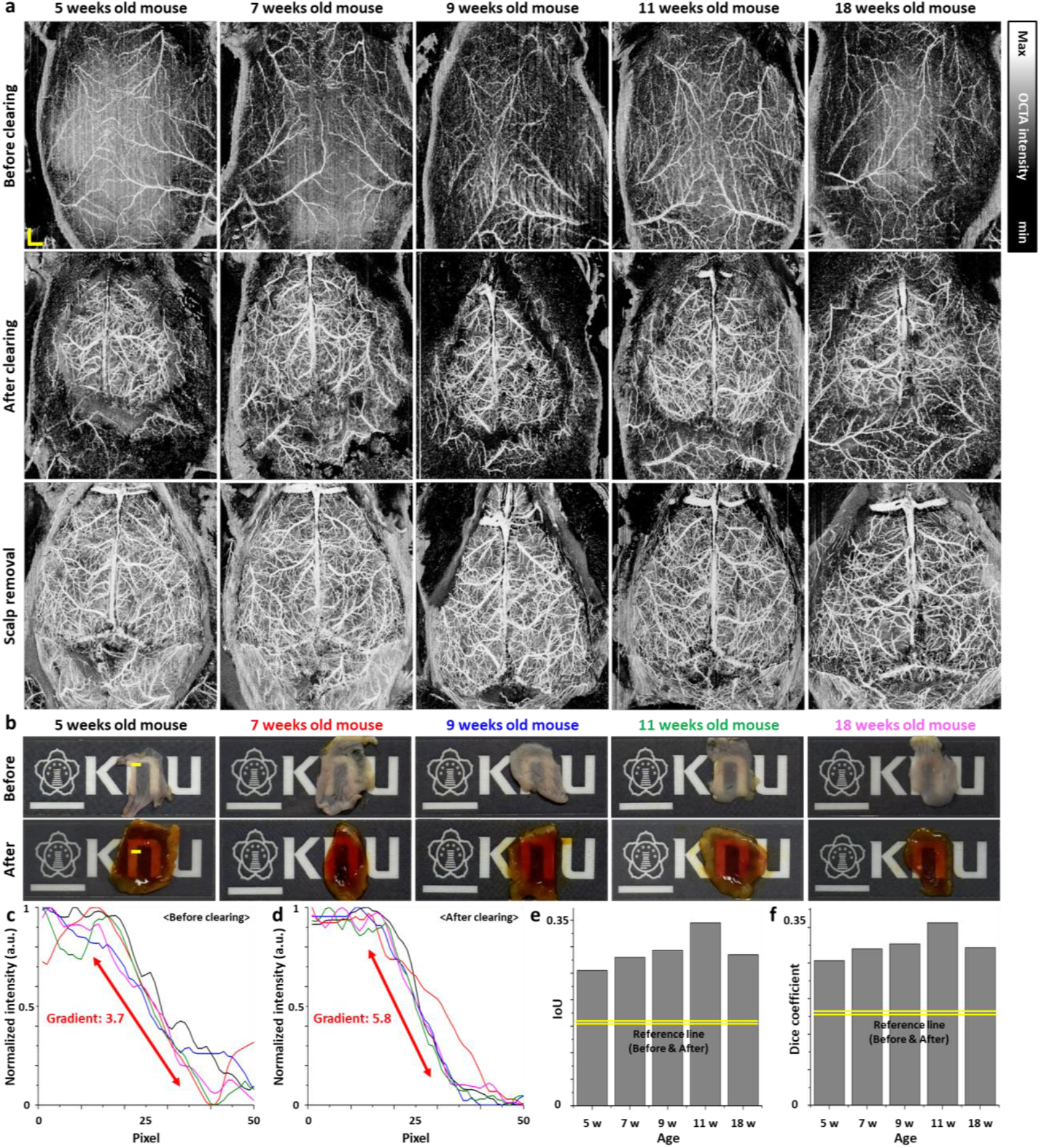
Validation results across mice of different ages from 5 weeks to 18 weeks. **a** OCTA MIP images of different ages acquired at before optical clearing, after optical clearing, and scalp removal. **b** Photographs illustrating before and after optical clearing results at different ages of mice. **c** Line profile of the mouse scalp before optical clearing indicated with yellow line at **b**. The plot line colors correspond to the age of mouse colored in panel **b. d** Line profile of the mouse scalp after optical indicated with yellow line at **b. e** Calculated IoU values of different ages using magnified images at after optical clearing/scalp removal. **f** Calculated Dice coefficient values of different ages using magnified images at after optical clearing/scalp removal. IoU, intersection over union; OCTA, optical coherence tomography angiography. Scale bar at **a**: 1 mm. Scale bar at **b**: 1 cm.

## Discussion

In this work, we introduced a tartrazine-based intact-scalp whole mouse brain OCTA imaging protocol. By leveraging tartrazine-based optical clearing, our approach mitigates the optical barrier imposed by the scalp and enables intact-scalp OCTA visualization of mouse brain vasculature without scalp excision. We first established feasibility in the NIR-II spectral regime of our OCTA system and demonstrated region-specific clearing, in which intracranial vessels emerged selectively within the tartrazine-treated ROI while untreated areas retained superficial scalp-vascular features. This transition from scalp-dominant signals to depth-resolved cerebral vasculature was supported by depth-encoded projections and cross-sectional OCT/OCTA, which showed increased signal recovery at depth and an extended vessel-detection range after clearing. Quantitatively, the clearing-induced change in vascular identity was captured using overlap metrics, with higher IoU and Dice values outside the ROI (where scalp vessels were preserved) and reduced similarity within the ROI (where brain vasculature replaced the pre-clearing scalp pattern). We further confirmed repeatability across three independent mice and validated anatomical specificity by showing close agreement between post-clearing intact-scalp OCTA and OCTA acquired after scalp removal. Systematic concentration screening identified 0.6 M tartrazine as an optimal condition that provided a sufficiently large cleared field for whole-brain scanning while avoiding the rapid solidification that limited higher concentrations. Finally, we demonstrated applicability across a broad age range (5-18 weeks), indicating that tartrazine-assisted intact-scalp OCTA can provide a practical and generalizable route to non-invasive structural mapping of mouse brain vasculature.

Beyond enabling intact-scalp OCTA, our results highlight two practical features that are particularly relevant for routine preclinical imaging. First, the localized nature of tartrazine application provides an internal control within the same animal, as out-of-ROI regions largely retain superficial scalp-vascular patterns while the ROI undergoes a clear transition toward intracranial vascular morphology. This spatial specificity supports the interpretation that the observed contrast change is driven by optical clearing rather than global acquisition variability. Second, solution handling emerged as a key determinant of whole-brain imaging applicability. Although higher molar concentrations improved transparency, rapid solidification at 0.7–0.8 M reduced the effective imaging window and introduced occluded regions during large-area scanning, introducing occluded regions during large-area scanning, indicating that practical handling constraints (e.g., solidification over time) must be considered alongside clearing efficacy when selecting an optimal formulation for intact-scalp volumetric OCTA. Mechanistically, the robustness of clearing across ages–despite expected age-related increases in scalp thickness and scattering–suggests that tartrazine primarily improves trans-scalp light delivery by reducing scattering through refractive-index matching within the scalp microstructure, consistent with dye-mediated refractive-index modulation described by the Kramers–Kronig framework^25,26^. Together, these considerations position tartrazine-based scalp clearing as a practical, controllable front-end strategy for OCTA, while also motivating future work to improve formulation stability (e.g., drying/gelation control)^33^ and to quantify how residual skull-induced scattering and aberrations may still limit deeper vascular sensitivity^34,35^.

The present results demonstrate that tartrazine-based optical clearing can provide a practical route to intact-scalp OCTA visualization of mouse brain vasculature by improving trans-scalp optical access in the NIR-II operating band of our system. Across experiments, intracranial vascular morphology emerged selectively within the tartrazine-treated region, whereas untreated areas retained superficial scalp-vascular features, and the post-clearing vascular maps showed close agreement with scalp-removed reference images, supporting the cerebral origin of the observed networks. Repeatability across independent animals, the identification of an operationally optimal concentration (0.6 M) for whole-brain scanning within a workable time window, and successful application across a broad age range (5–18 weeks) collectively indicate that the protocol can be deployed under typical preclinical imaging conditions without requiring surgical scalp excision. Looking forward, this capability may expand the feasibility of non-invasive, repeated structural angiography in longitudinal mouse studies where surgical cranial preparation is undesirable, including investigations of vascular remodeling, angiogenesis, and treatment-associated changes in vascular architecture.

However, there remains considerable room to further refine this technique. First, because optical clearing in our protocol requires massage after tartrazine application, the degree of transparency—and consequently the visibility of cerebral vessels—can vary depending on how effectively the scalp is massaged. This introduces potential operator-dependent variability that should be addressed through improved standardization. In addition, the extent of clearing may differ across individual animals depending on their physiological condition, indicating the need for additional protocol refinement and objective criteria for application. Finally, because the scalp is not rendered perfectly transparent and residual scattering persists, the resulting OCTA images may exhibit discrepancies from the true vascular morphology, for example by making vessels appear artificially thicker. To mitigate such effects, it will be important to quantitatively assess the degree of clearing and incorporate an image post-processing or correction step that accounts for residual scattering.

## Methods

### Optical coherence tomography angiography system configuration

OCTA imaging of tartrazine-based intact-scalp mouse brain was performed with a customized tomographic system. A MEMS-VCSEL swept source (SL134051, Thorlabs, USA), operating at a 400 kHz sweep rate with a 1300 nm central wavelength and 100 nm bandwidth, was employed to achieve a 3 mm imaging depth. The output of the source was split by a 90:10 fiber coupler (TW1300R2A2, Thorlabs, USA), with the light subsequently routed through a fiber optic circulator (CIR-1310-50-APC, Thorlabs, USA) to the reference and sample arms. The reference arm consisted of a collimator (F260APC-C, Thorlabs, USA), an achromatic lens (AC254-030-C, Thorlabs, USA), and a silver mirror (PF10-03-P01, Thorlabs, USA). The sample arm incorporated a collimator (F260APC-C, Thorlabs, USA), a two-dimensional galvanometer scanner (GVS102, Thorlabs, USA), and a scan lens (LSM04, Thorlabs, USA) featuring an effective focal length of 54 mm. The interference signal was collected by a 50:50 coupler (TW1300R5A2, Thorlabs, USA), detected via a balanced photodetector (PDB480C-AC, Thorlabs, USA), and digitized by a digitizer (ATS9373, Alazar Technologies Inc., Canada).

### Optical coherence tomography angiography processing pipeline

For OCTA acquisition, we used an A-scan rate of 200 kHz and acquired six repeated B-scans at each lateral position along the slow axis, with each B-scan consisting of 2,000 A-lines sampled along the fast axis. This procedure was repeated across 2,000 positions along the slow axis to form a volumetric dataset. Raw OCT interferograms were processed using a standard OCT reconstruction pipeline, including background subtraction and fast Fourier transformation to obtain depth-resolved OCT signals. In terms of the k-linearization, since we used k-clock from the source as a trigger, therefore, the obtained raw graph was not required for linearization. To extract microvascular information, we employed an amplitude-decorrelation OCTA algorithm^14^. To reduce noise-related bias in decorrelation estimation, we applied an intensity threshold to exclude low-SNR pixels and subsequently performed median filtering on the decorrelation images.

### Image quality evaluation metrics

To quantitatively compare whether two OCTA datasets represented the same vascular patterns within a given field of view (e.g., before vs after clearing, or after clearing vs scalp removal), we computed binary overlap metrics (IoU and Dice coefficient) from vessel-segmented OCTA maximum-intensity-projection (MIP) images using a standardized MATLAB pipeline implemented with the Image Processing Toolbox. OCTA MIP images were exported as PNG files and robustly converted to grayscale intensity images in [0, 1] regardless of input type (indexed, RGB, or grayscale). If an alpha channel was present, transparent pixels were masked to zero. Image pairs were cropped to a common overlapping size to ensure pixel-wise correspondence. To avoid per-image contrast stretching from artificially inflating similarity, both images in a pair were normalized using a shared minimum and maximum intensity computed from the concatenated pixel values of the two images (common min–max normalization). Vessel structures were then enhanced using a multiscale Frangi-like vesselness procedure by applying Gaussian smoothing across multiple scales (*σ*=1–5 pixels) followed by a vesselness response, and the maximum response across scales was retained and normalized to [0, 1]. Binary vessel masks were generated by thresholding the vesselness images with an automatic threshold selection scheme designed to control segmentation density. After thresholding, small isolated components were removed using an area filter with a minimum size of 80 pixels.

Binary overlap between the two vessel masks was quantified using IoU and Dice coefficient. Let M_1_ and M_2_ denote binary vessel masks (vessel pixels = 1, background = 0) computed from the two images being compared within the same region. IoU (Jaccard index) and Dice were defined as

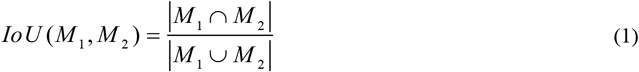

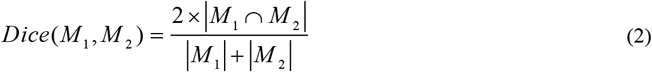

where |·| indicates the number of vessel pixels. Equivalently, in terms of pixel-wise true positives (TP), false positives (FP), and false negatives (FN),

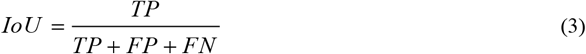

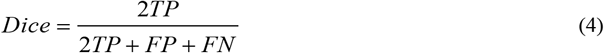

IoU and Dice were computed for each image pair within the corresponding analysis region, and higher values indicate greater spatial agreement of the segmented vascular patterns between the two conditions.

### Tartrazine-enabled mouse brain OCTA imaging protocol

A 0.6 M tartrazine optical clearing formulation was prepared by adding 3 g of tartrazine powder to an empty vial and dispensing 7.0 mL of Milli-Q water. The mixture was vortexed until homogeneous, capped, and incubated in a preheated 80 °C oven for 10 min. After cooling for 5 min, exactly 5 g of the tartrazine solution was transferred to a new vial, supplemented with 30 mg of low-melting-temperature SeaPlaque agarose, loosely covered, and heated at 80 °C for 1 h until the agarose was fully dissolved. The vial was then capped and stored until use. For in-vivo imaging, C57BL/6 mice were prepared by shaving the scalp and removing residual hair using depilatory cream for 3 min, followed by gentle wiping with water-moistened gauze. Baseline OCTA images were acquired before clearing. The tartrazine formulation was applied dropwise onto the shaved scalp using a disposable dropper while the scalp surface was gently massaged with a cotton swab for 7 min. Excess solution was removed from the surrounding area using gauze, and OCTA images were acquired after clearing. Finally, the optically cleared scalp region was excised using surgical scissors, and OCTA images were acquired again under the scalp-removed condition as a reference for cerebral vasculature visualization. An identical optical clearing protocol was applied across all age groups and tartrazine molar concentrations. All experimental procedure of this study was conducted according to the guidelines of the Declaration of Helsinki and approved by the Institutional Animal Care and Use Committee of Kyungpook National University (KNU-2025-0845).

### UV-VIS-NIR spectroscopy

Optical transparency of the dye solutions in the 1.2–1.4 µm wavelength range was measured using a UV–VIS–NIR spectrophotometer (SolidSpec-3700i, Shimadzu). Transmittance measurements were performed with a standard quartz cuvette having a 1 mm optical path length. The reference compartment was left empty, and all measurements were conducted relative to air.

### Ellipsometry measurements of the RI of dye solutions

Refractive indices of the dye solutions listed in the table were measured using an ellipsometer (Alpha 2.0, J.A. Woollam). For each measurement, 4 mL of solution was poured into a disposable base mold (37 mm × 24 mm × 5 mm; 62352-37, Electron Microscopy Sciences), fully covering the cavity bottom to form a flat, reflective air–liquid interface. The ellipsometer operates over a wavelength range of 0.3–1.0 µm; therefore, refractive index values in the 1.2–1.4 µm range were obtained through model-based extrapolation. Specifically, the experimentally measured refractive index data were fitted using the Cauchy dispersion model, and the fitted parameters were subsequently used to extend the refractive index values to the desired near-infrared wavelength range. This extrapolation and analysis were performed using MATLAB (Supplementary Fig. 2).

## Data availability

Raw experimental data supporting the findings of this study are available from the corresponding author upon request.

## Acknowledgements

This work was supported by the Institute of Information & Communications Technology Planning & Evaluation (IITP)-Innovative Human Resource Development for Local Intellectualization program grant funded by the Korea government (MSIT)(IITP-2025-RS-2022-00156389).

