## Supplementary information for "Beyond the skin barrier: optical clearing enables non-invasive cortex-wide optical coherence angiography in mice in-vivo"

**Supplementary Figure 1.** Measured refractive indices of the tartrazine solutions with different molar concentration in the 0.6 to 1.0 μm wavelength range.

**Supplementary Figure 2.** Measured refractive index and extrapolation for 0.6 M tartrazine solution using the Cauchy dispersion model.

**Supplementary Figure 3.** Color-coded depth mapping of mouse scalp vasculature before and after clearing.

**Supplementary Figure 4.** Color-coded depth mapping of mouse scalp vasculature before and after optical clearing with 0.3 M-0.8 M tartrazine.

**Supplementary Figure 5.** Magnified OCTA images at before optical clearing, after optical clearing, and scalp removal with different molar concentrations.

**Supplementary Figure 6.** Photographs of tartrazine solidification over time at different concentrations of 0.3 M-0.8 M.

**Supplementary Figure 7.** Effect of the solidification in OCTA images at high concentration tartrazine (0.7 M and 0.8 M).

**Supplementary Figure 8.** Color-coded depth mapping of mouse scalp vasculature before optical clearing and after optical clearing at different ages.

**Supplementary Figure 9.** Magnified OCTA images before clearing, after clearing, and scalp removal at different ages.

**Supplementary Table 1.** Calculated mass of tartrazine for 10 ml DI water from 0.3 M to 0.8 M.

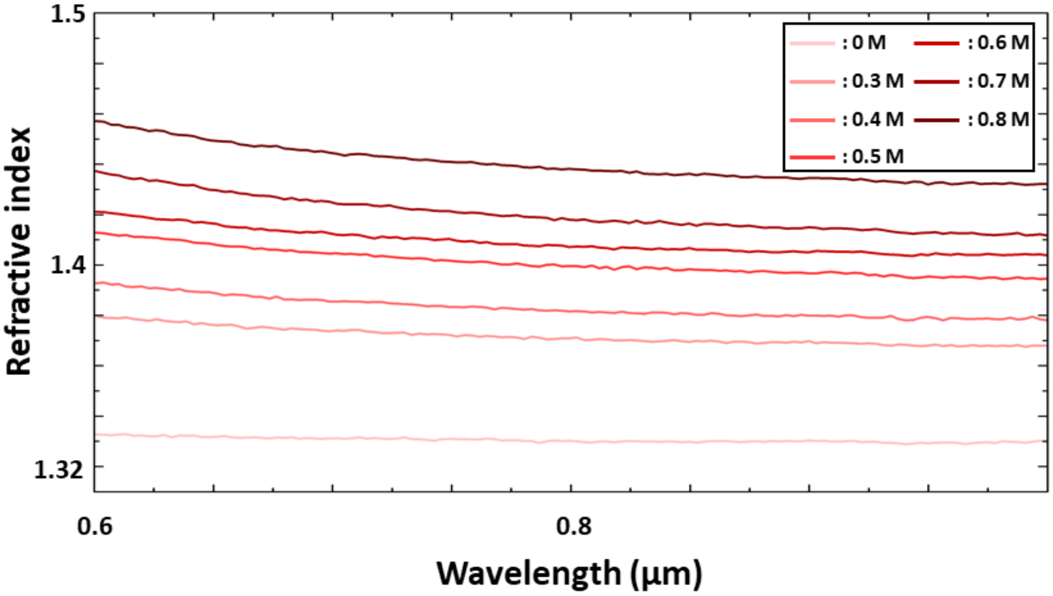


**Supplementary Figure 1. Measured refractive indices of the tartrazine solutions with different molar concentration in the 0.6 to 1.0 μm wavelength range.**

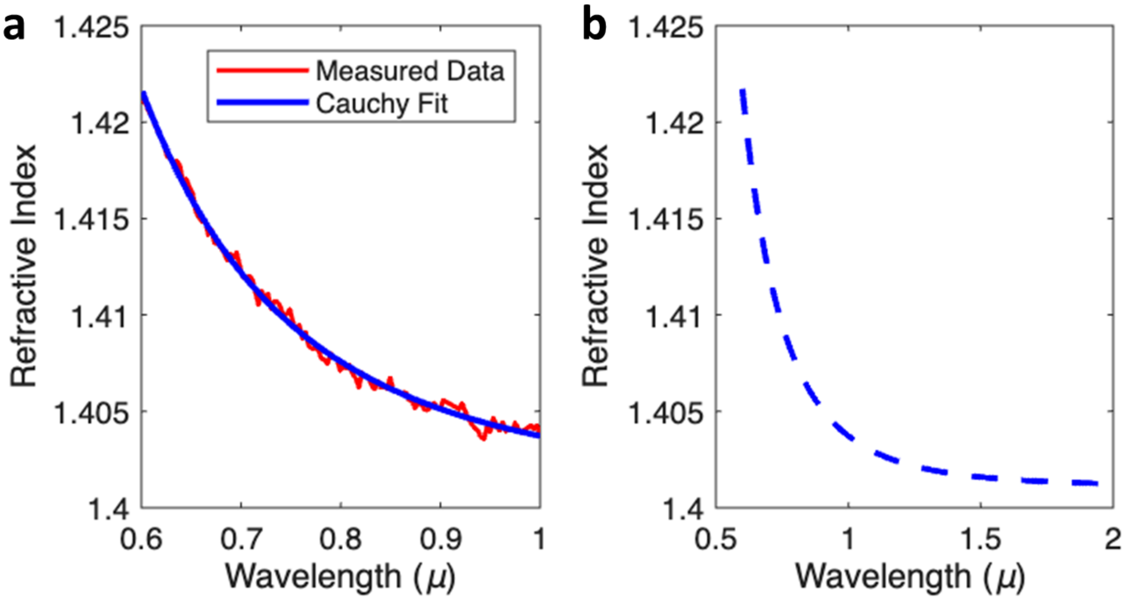


**Supplementary Figure 2. Measured refractive index and extrapolation for 0.6 M tartrazine solution using the Cauchy dispersion model. a** Experimental refractive index values measured using an ellipsometer in the wavelength range of 0.6-1.0 μm. **b** Extended refractive index spectrum extrapolated up to 2 μm using the Cauchy dispersion model based on the experimental data.

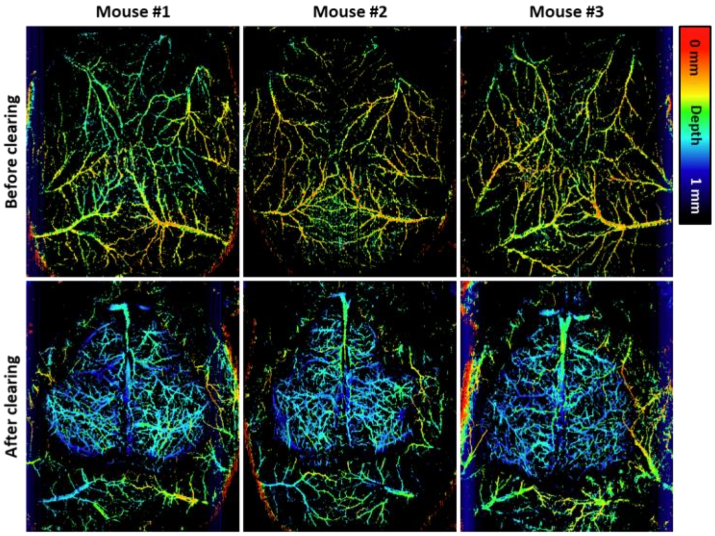


**Supplementary Figure 3. Color-coded depth mapping of mouse scalp vasculature before and after clearing.**

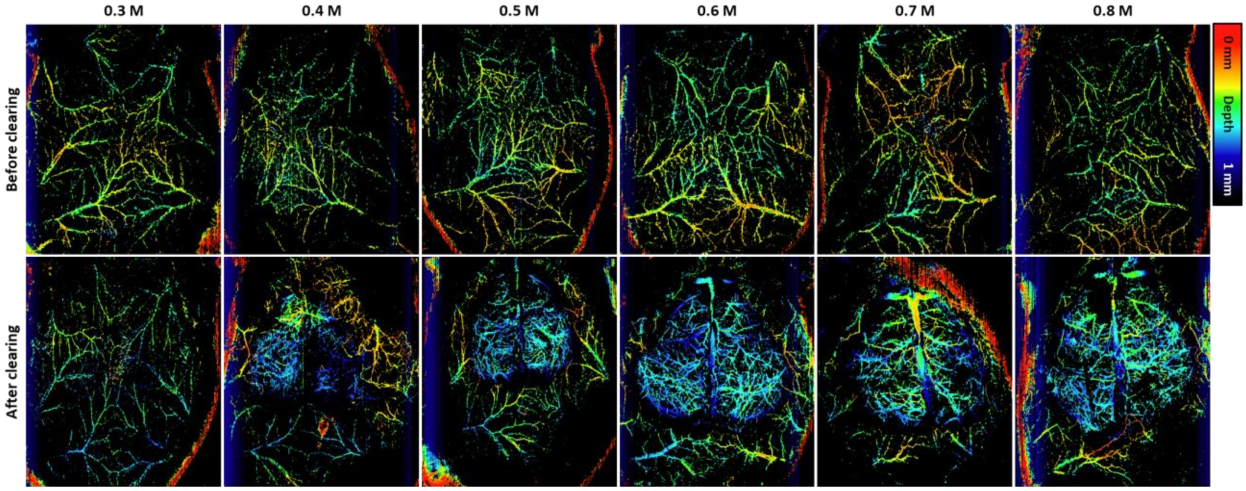


**Supplementary Figure 4. Color-coded depth mapping of mouse scalp vasculature before and after optical clearing with 0.3 M-0.8 M tartrazine.**

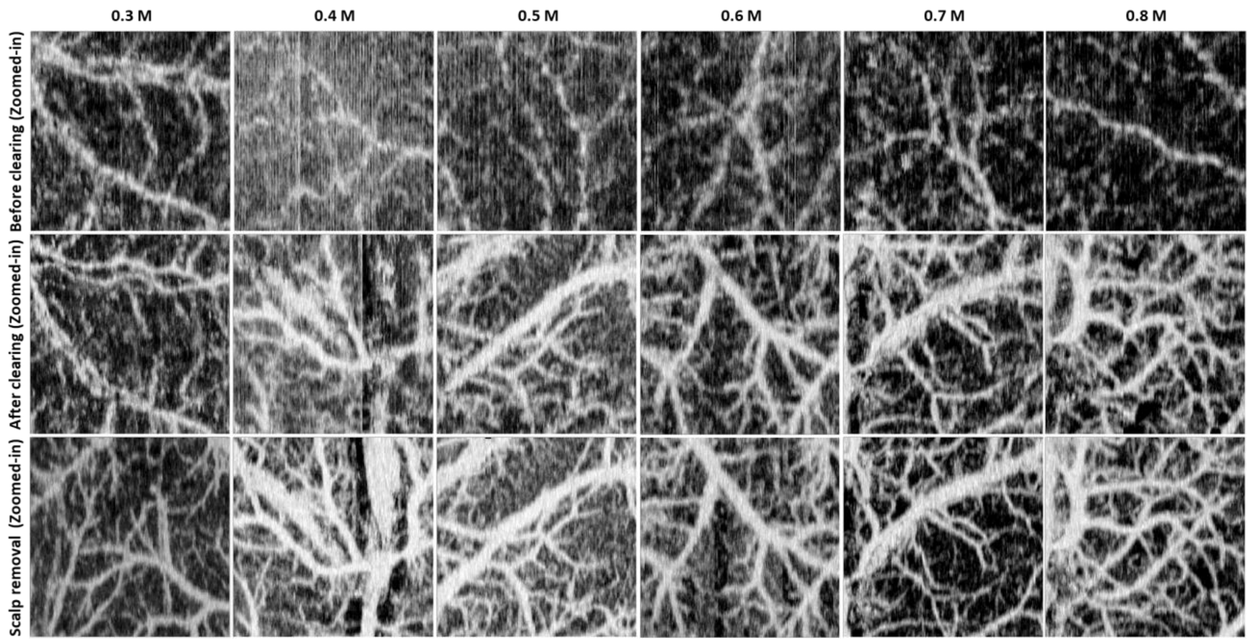


**Supplementary Figure 5. Magnified OCTA images at before optical clearing, after optical clearing, and scalp removal with different molar concentrations.**

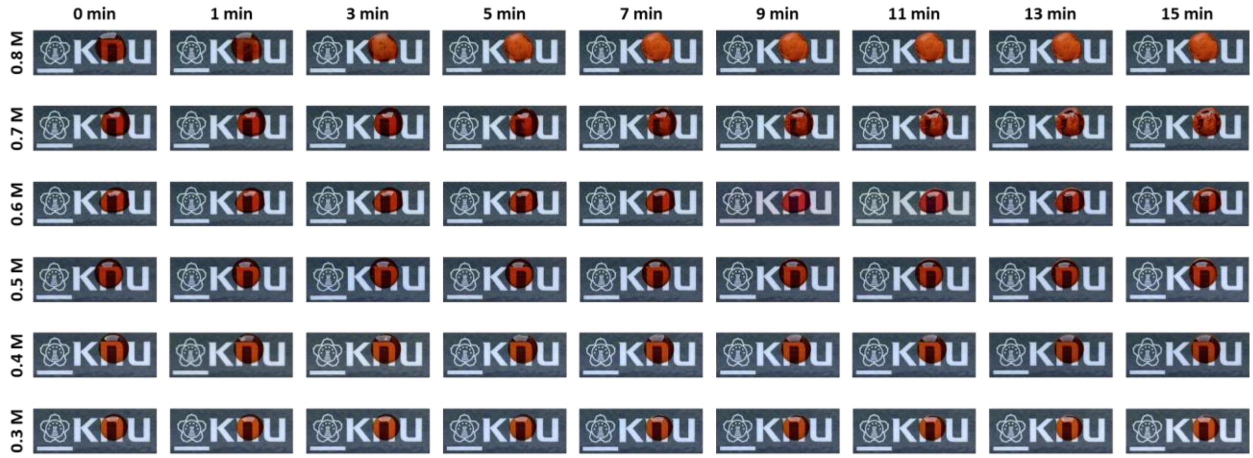


**Supplementary Figure 6. Photographs of tartrazine solidification over time at different concentrations of 0.3 M-0.8 M.**

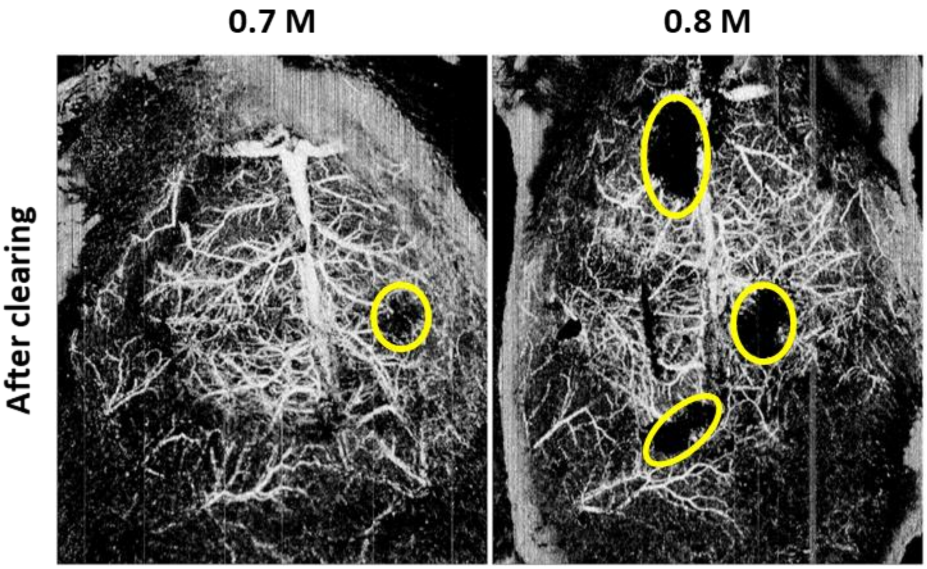


**Supplementary Figure 7. Effect of the solidification in OCTA images at high concentration tartrazine (0.7 M and 0.8 M).**

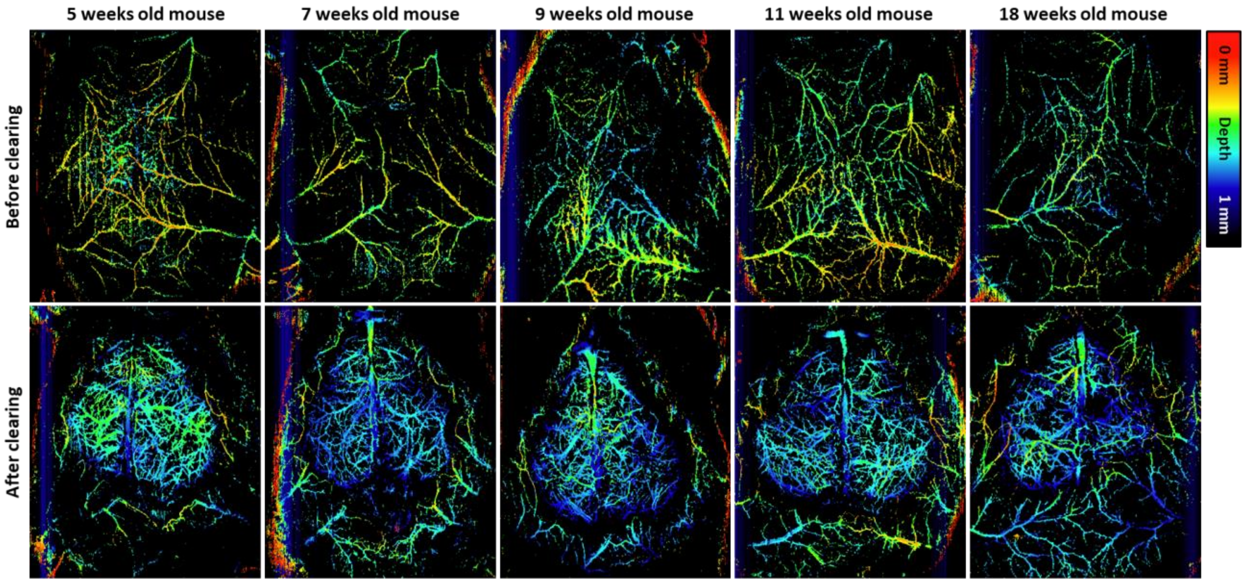


**Supplementary Figure 8. Color-coded depth mapping of mouse scalp vasculature before optical clearing and after optical clearing at different ages.**

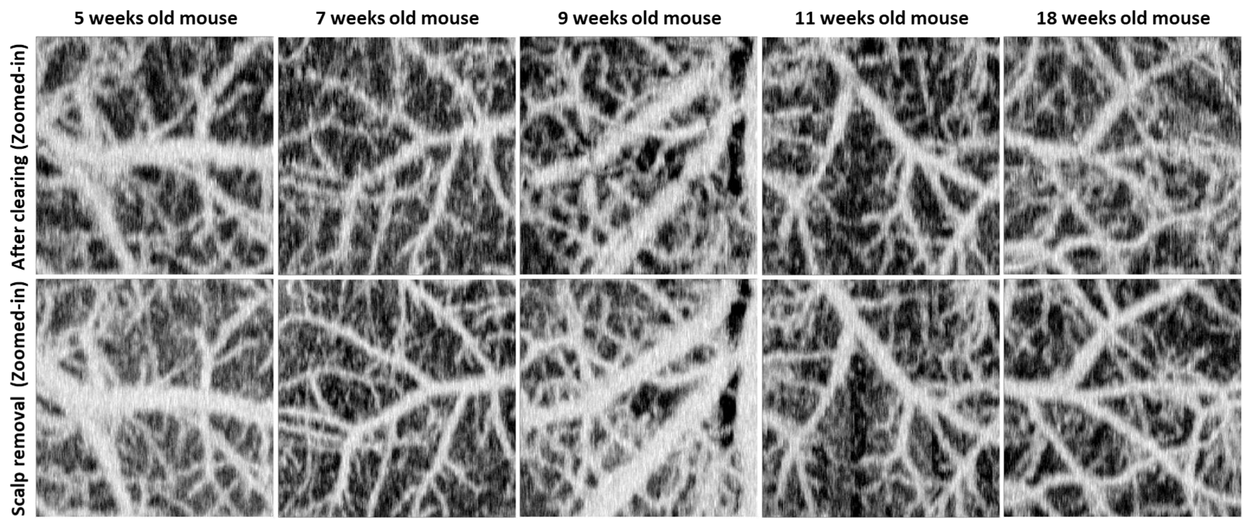


**Supplementary Figure 9. Magnified OCTA images before clearing, after clearing, and scalp removal at different ages.**

**Supplementary Table 1. Calculated mass of tartrazine for 10 ml DI water from 0.3 M to 0.8 M.**

| S.N. | Tartrazine concentration | Tartrazine mass dissolved in 10ml DI Water |
|---|---|---|
| 1 | 0.3 M | 1748.2 mg |
| 2 | 0.4 M | 2403.6 mg |
| 3 | 0.5 M | 3101.0 mg |
| 4 | 0.6 M | 3484.9 mg |
| 5 | 0.7 M | 4639.7 mg |
| 6 | 0.8 M | 5491.1 mg |
